# Mapping Focal and Generalized Effects of Common Genetic Variants on Human Brain Structure

**DOI:** 10.64898/2026.07.15.738539

**Authors:** Emma J. Gleave, Luis M. García-Marín, Zuriel Ceja, Miguel E. Rentería, Tamoghna Chattopadhyay, Christian Gaser, Priya Rajagopalan, Paul M. Thompson

**Affiliations:** Imaging Genetics Center, Mark and Mary Stevens Neuroimaging and Informatics Institute, Keck School of Medicine, University of Southern California, CA, USA; Brain and Mental Health Program, QIMR Berghofer, Brisbane, Queensland, Australia; Jena University Hospital, Jena, Germany; Department of Radiology, Los Angeles General Medical Center, Los Angeles, CA, USA

**Keywords:** Voxel-based Morphometry, Multivariate imaging genetics, GWAS, PGS, Structural MRI

## Abstract

Genome-wide association studies (GWAS) have advanced the quest to understand how specific genetic variants influence human brain structure and function. Recent work has identified hundreds of common variants associated with subcortical brain volumes, sparking interest in how these genetic markers overlap across brain networks. While this can be estimated by hierarchical clustering of the genetic correlation matrix to identify modular patterns of shared architecture, no brain-wide maps of these effects are available. To address this, we computed polygenic scores (PGS) from loci associated with ten brain volume regions of interest (ROIs): nine major subcortical structures and intracranial volume, with each locus weighted by its association with regional volume. In an independent sample from the discovery GWAS, we performed large-scale segmentation of 3D volumetric T1-weighted MRI scans using voxel-based morphometry (VBM) to map 3D profile of regions where gray matter volume (GMV) was associated with each PGS. We found statistically significant, localized effects for PGS defined for the amygdala, thalamus, and basal ganglia, but PGS for brainstem volume was associated with widespread differences throughout the brain. These brain-wide maps reveal patterns consistent with both localized and distributed genetic influences, offering a novel approach to interpret the genomic architecture of brain structure.

## I. Introduction

There is great interest in mapping how genes shape brain development, aging, and disease. One central question in uncovering this process is whether specific genetic variants influence these traits of development or disease in smaller, region-specific patterns rather than in widespread effects across multiple brain regions. Distinguishing between global vs localized effects is critical for linking genetic architecture to brain processes, whether investigating traits related to neurodevelopmental trajectories, age-related changes, or predisposition to neurological or neuropsychiatric disorders. Such traits are frequently examined utilizing genome-wide association studies (GWAS): large-scale studies in which the entire genome is scanned for common genetic variants associated with those specific traits or phenotypes. In the context of brain imaging, GWAS can identify common genetic variants that influence the morphometry or connectivity of different brain regions, thus allowing distinction between genetic effects that are broadly distributed across the brain versus those that are localized to specific regions [1,2,3], including subcortical [4,5]. A recent GWAS in over 75,000 individuals scanned with MRI from Garcia-Marin et al. (2024) identified more than 500 common variants associated with subcortical brain volumes [6], sparking interest in how strongly these genetic markers overlap across different brain networks. However, while this can be estimated by hierarchical clustering of the genetic correlation matrix to identify ‘genetic modules’ (*i*.*e*., clusters of ROIs with high genetic correlation) [7,8,9] no brain-wide maps of these effects are available. As maps of brain structure and function are inherently multivariate, and such variation is highly polygenic (*i*.*e*., numerous genetic variants simultaneously influence multiple regions), alternative imaging genetic methods beyond univariate GWAS and hierarchical clustering are required to uncover the polygenic architecture underlying brain-related phenotypes further.

Multivariate imaging genetic approaches typically combine multiple functional or anatomical imaging variables with many genetic variables, to uncover coordinated or distributed patterns linking genetic variants to brain structure and function [10,11,12], Due to their higher dimensionality, multivariate methods may identify broader polygenic architecture and enable a more robust assessment of biologically meaningful patterns [13,14]. Various approaches to multivariate imaging genetics exist. Parallel independent component analysis (pICA) is commonly used in this context, jointly decomposing imaging and genetic data into independent components whose subject-level loadings are maximally associated across modalities. These linked components capture coordinated genetic influences on distributed brain networks. As such, pICA can reveal linked components of brain structure and genetic variation that would be missed by univariate analyses, highlighting shared polygenic influences on neural systems [15,16]. Other relevant approaches include canonical correlation analysis (CCA), which formulates an optimization problem to identify linear combinations of genetic variants and imaging features whose projected scores are maximally correlated across subjects. This can then be used to identify multivariate relationships between genomes and brain structure. Shen et al. (2020) showed that deep sparse CCA could detect distributed polygenic effects on brain connectivity, a connection not likely to be found using more traditional methods, improving interpretability and prediction of complex traits [17]. Zhan et al. (2025) proposed a transformer-based framework for imaging genetics that uses attention mechanisms to capture nonlinear, high-dimensional relationships between genetic variation and distributed brain phenotypes, showing improved performance over conventional linear multivariate approaches [18].

Here, we set out to ascertain whether genetic loci identified through conventional GWAS exert their effects in a region-specific manner or more broadly across the brain. By focusing on loci robustly associated with subcortical and intracranial volumes, we aimed to map the spatial distribution of polygenic effects, linking large-scale genomic findings to anatomically interpretable patterns of gray matter variation. We computed polygenic scores (PGS)[19] from loci associated with ten brain volume regions of interest (ROIs): nine major subcortical structures and intracranial volume (ICV). We used voxel-based morphometry (VBM) to map brain signatures of regional gray matter volume (GMV) relative to each brain volume ROI PGS after accounting for factors such as age and sex. Additionally, by generating Quantile-Quantile (QQ) plots to visualize statistical significance values, we were able to evaluate an overall measure of the strength of association. This mass univariate testing approach combines imaging, in which VBM allows for spatial anatomic mapping, with genetics via PGS to consolidate effects across multiple loci. Thus, it integrates within multivariate imaging genetics frameworks by incorporating voxelwise PGS maps as high-dimensional imaging features in statistical models that jointly relate patterns of genetic variation to distributed brain structure. Our approach links large-scale GWAS of brain volumes to spatially resolved, anatomically interpretable representations of polygenic effects, enabling systematic comparison of shared and distinct genetic influences on brain organization.

## II. Data and Methods

### A. Participants and MRI Acquisition

Our primary study cohort for the PGS-VBM analyses included a subset of UK Biobank (UKB) data. The UKB is a longitudinal prospective population study of over 500,000 middle-aged to older adults, featuring a vast database of clinical health and demographic measures collected from multiple sites across the UK [20,21]. In addition to the comprehensive phenotypic data available, including extensive imaging data such as T1-weighted (T1w) brain MRI, the UKB also contains a rich repository of genetic data. Notably, it provides pre-computed genetic principal components (PCs) derived from principal component analysis (PCA) of the genome-wide genotype data. In this study, we included the first four PCs, which summarize major axes of population structure related to ancestry, as covariates [22,23].

To ensure sample independence, we selected UKB participants who had not been included in our initial discovery GWAS from Garcia-Marin et al. (2024) [6]. This was done to avoid circularity and overfitting of locus weighting and thereby provide an unbiased assessment of the relationship between genetic predisposition for regional brain volume and voxelwise gray matter morphometry. Criteria for inclusion in this subset sample required the availability of imaging data (a baseline T1w 3D volumetric brain MRI), genome-wide genetic data (including derived scores for the first four PCs reflecting ancestry and population structure, as is common in genomic analysis), and simple demographics (age and sex). Participants were excluded from analyses if these were missing or failed quality control checks, such as poor image resolution or scan motion. The final set of 2,830 participants (age: 68.8 ± 8.1 years; 51.0% female) from across four scan sites was used as an independent sample for VBM analyses to map the 3D profile of brain regions where GMV was associated with each PGS. We note that the original sample in which the genomic variants were discovered was much larger, so the goal of the current work was to map their effects with greater spatial precision, in an independent, non-overlapping sample not used in the original discovery GWAS.

### B. Genetic Data processing

We evaluated ten regional brain subvolume regions of interest (ROIs). These included 9 distinct subcortical structures: the amygdala, caudate nucleus, hippocampus, nucleus accumbens, globus pallidus, putamen, thalamus, ventral diencephalon, and brainstem, in addition to an overall measure of intracranial volume (ICV). PGS were derived for each ROI, where each locus was weighted by its strength of association with regional volume, in the original GWAS. Specifically, for each individual *i* and ROI (*r*), the PGS was computed as:

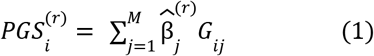

where *j* indexes genetic loci represented by single-nucleotide polymorphisms (SNPs), *M* denotes the number of SNPs included in the score, *G*_*ij*_ is the number of effect alleles (0, 1, or 2) carried by individual *i* at SNP *j*, and 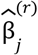 is the estimated effect size 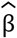 of SNP *j* on ROI *r*. To derive the PGS, SBayesR was run [24], within the Genome-wide Complex Trait Bayesian analysis (GCTB v2.0) software tool [25]. SBayesR jointly models all SNPs while accounting for linkage disequilibrium and the sparse, polygenic nature of brain volume traits, producing shrinkage-adjusted effect estimates that optimize predictive accuracy. Each PGS thus represents an individual’s aggregate genetic predisposition for variation in a specific ROI.

### C. Image Data processing

To map the anatomical expression of these polygenic influences, we used voxel-based morphometry (VBM) to measure gray matter volume (GMV) at each voxel across the brain [26]. Image processing and gray matter VBM segmentation of the T1w brain MRI scans was implemented using the ENIGMA Computational Anatomy Toolbox (**Figure 1**; CAT12 [https://neuro-jena.github.io/enigma-cat12/]). [27] Implemented within standard Statistical Parametric Mapping (SPM) MATLAB software [28]. This pipeline, which is widely used, performs a large-scale voxel-wise segmentation of the 3D T1w brain MRI data after running standard image preprocessing steps, such as denoising, bias correction, and spatial registration. VBM outputs from the CAT12 pipeline include a 3D whole-brain image mask of the participant’s segmented gray matter voxels and another for white matter, as well as tabulated volumetric estimations for overall gray matter, white matter, CSF, and total intracranial volume (TIV). During registration, the gray matter tissue class masks were spatially normalized and modulated by the Jacobian determinants of the deformation fields, preserving local gray matter volume prior to smoothing and statistical analysis; following CAT12 processing, the gray matter brain masks were then smoothed with a 3D Gaussian spatial smoothing kernel. Typical smoothing kernel sizes range from 2-mm or 4-mm up to 12-mm depending on the scope and ROI being analyzed [27,29]. However, 8-mm is the default size smoothing kernel in SPM and therefore commonly employed in VBM processing, and was selected here in our study as a midrange value to preserve adequate anatomical regional specificity while still maintaining sensitivity of results [30,31]. This balance was important for our analyses, given the range of size of subcortical structures in our study, while protecting against false positive results and inflated effect sizes that can arise on unsmoothed or lower-kernel smoothed data [30,32-34].

**Figure 1.**
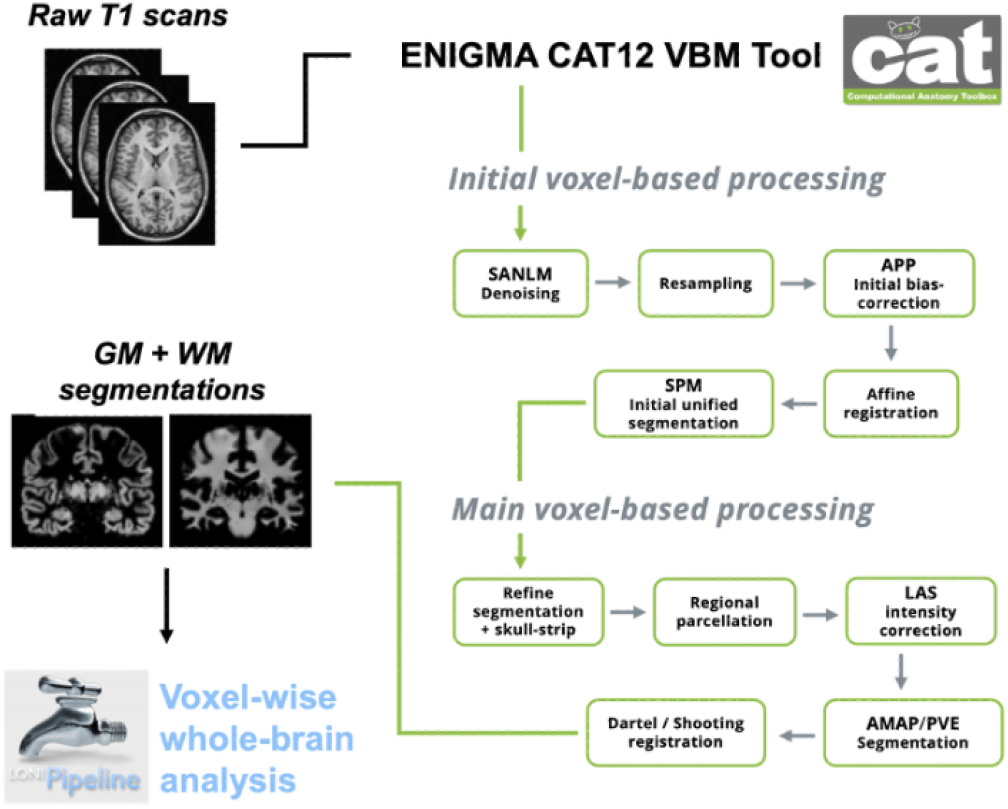
VBM Pipeline and Analysis Workflow. The ENIGMA CAT12 VBM pipeline (as used by the ENIGMA Consortium) processes raw 3D T1w MRI scans with a first module including a spatial adaptive non-local means (SANLM) denoising filter, bias correction, segmentation, and registration, followed by a second module for spatial registration including skull-stripping, regional parcellation, intensity transformations and refinement of brain tissue classification and volume estimations for voxel-wise analysis. LAS (Local Adaptive Segmentation) improves local tissue intensity normalization, AMAP (Adaptive Maximum A Posteriori) segmentation provides robust tissue classification without relying on population priors, and PVE (Partial Volume Estimation) addresses voxels containing mixtures of different tissues, enhancing segmentation accuracy at boundaries.

### D. Statistical Analysis

The statistical relationship between voxel-wise brain GMV *y* and each ROI PGS was assessed using a linear mixed model (**Figure 1**; LONI pipeline [http://pipeline.loni.usc.edu] v.7.0.3; using R version 2.9.2)[35] as follows:

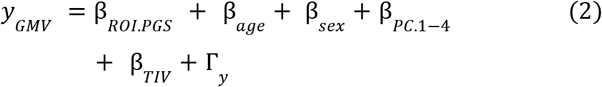

where covariates of age, sex, the first four PCs from the UKB PCA capturing genetic ancestry, and TIV (estimated via CAT12 VBM segmentation) were accounted for as fixed effects. Additionally, the scanner site was included as a random effect Γ_*y*_ to account for the potential effects of site-specific variance across UKB acquisition centers. Results were corrected for multiple comparisons using the standard false discovery rate (FDR) at 5% (q=0.05) [36]. Significance maps of GMV voxel-wise results for each ROI PGS analysis were generated using the FDR-corrected *p*-values from each analysis (**Figure 2**), and these threshold *p*-values for each analysis summarized in **Table 1**. Raw *p*-values from each ROI PGS analysis were also plotted alongside one another in a quantile-quantile plot (**Figure 3**), with FDR-correction thresholds shown as the horizontal cutoff line along the observed value axis. Since VBM analyses were run on the modulated normalized gray matter mask images, the reported associations reflect gray matter–specific volumetric variation. Subcortical structures such as the caudate nucleus and thalamus contain substantial white matter components, so deformation occurring within these regions may be underestimated, leading to diminished effect sizes in those regression models.

**Table 1.** Significance reporting for the results of each ROI PGS x GMV regression model.

|  |  |
| --- | --- |
| **ROI** | **FDR-critical $p$-value** |
| Amygdala | 0.0032 |
| Caudate Nucleus | 0.0012 |
| Hippocampus | 0.0028 |
| Nucleus Accumbens | 0.0014 |
| Globus Pallidus | 0.0013 |
| Putamen | 0.0031 |
| Thalamus | 0.0011 |
| Ventral Diencephalon | 0.0128 |
| Brainstem | 0.0239 |
| ICV (no TIV covariate) | 0.0498 |

**Fig. 2.**
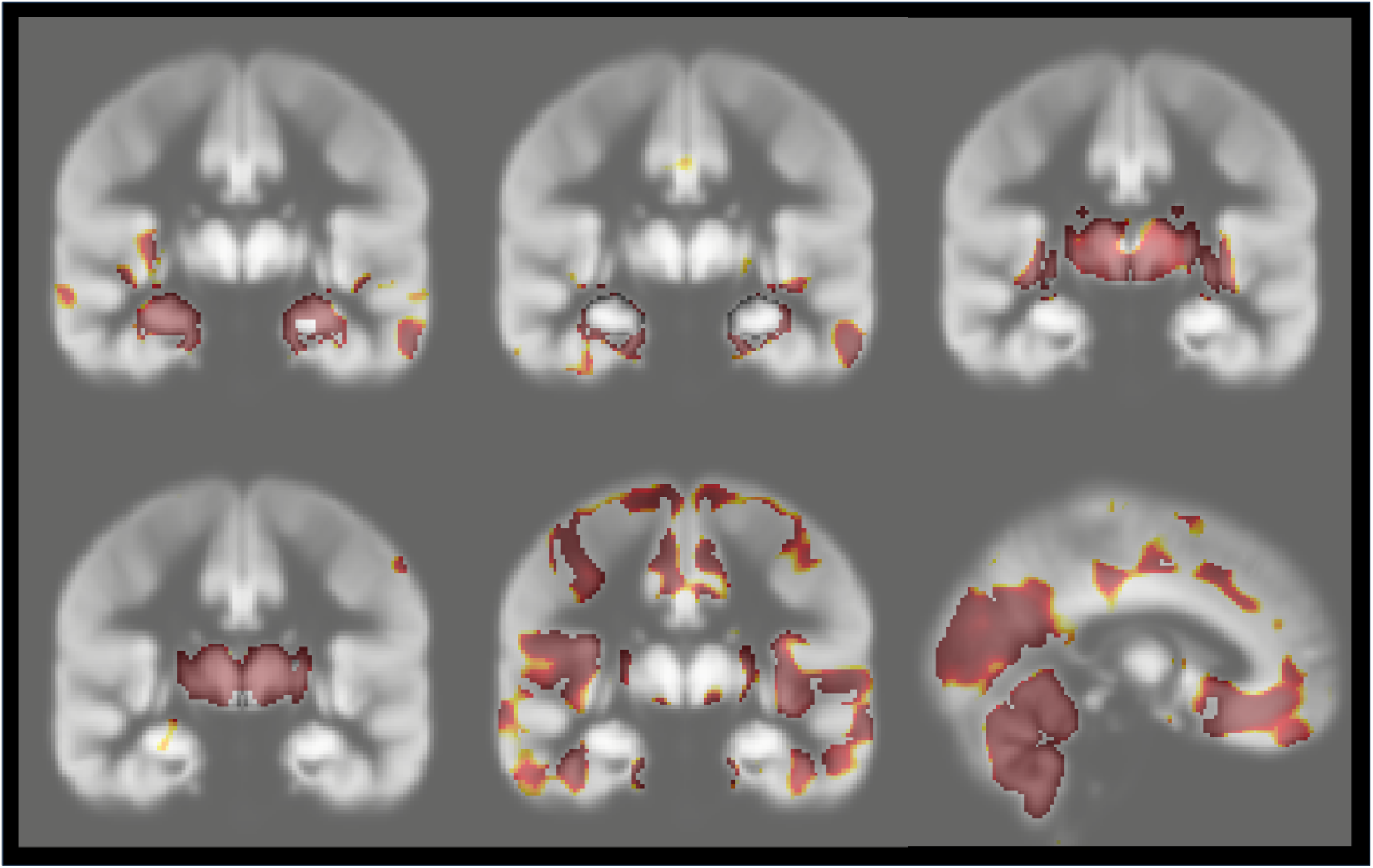
FDR-corrected (q=0.05) voxel-wise whole-brain significance maps of the association between polygenic score for brain ROI and GMV from linear mixed model analyses of 2,830 independent UKB participants: amygdala (*top left*; standard-FDR critical *p*-value=0.0032), hippocampus (*top center*; standard-FDR critical *p*-value=0.0028), putamen (*top right*; standard-FDR critical *p*-value=0.0031), thalamus (*bottom left*; standard-FDR critical *p*-value=0.0011), and the brainstem (*bottom center, bottom right*; standard-FDR critical *p*-value=0.0239).

**Fig. 3.**
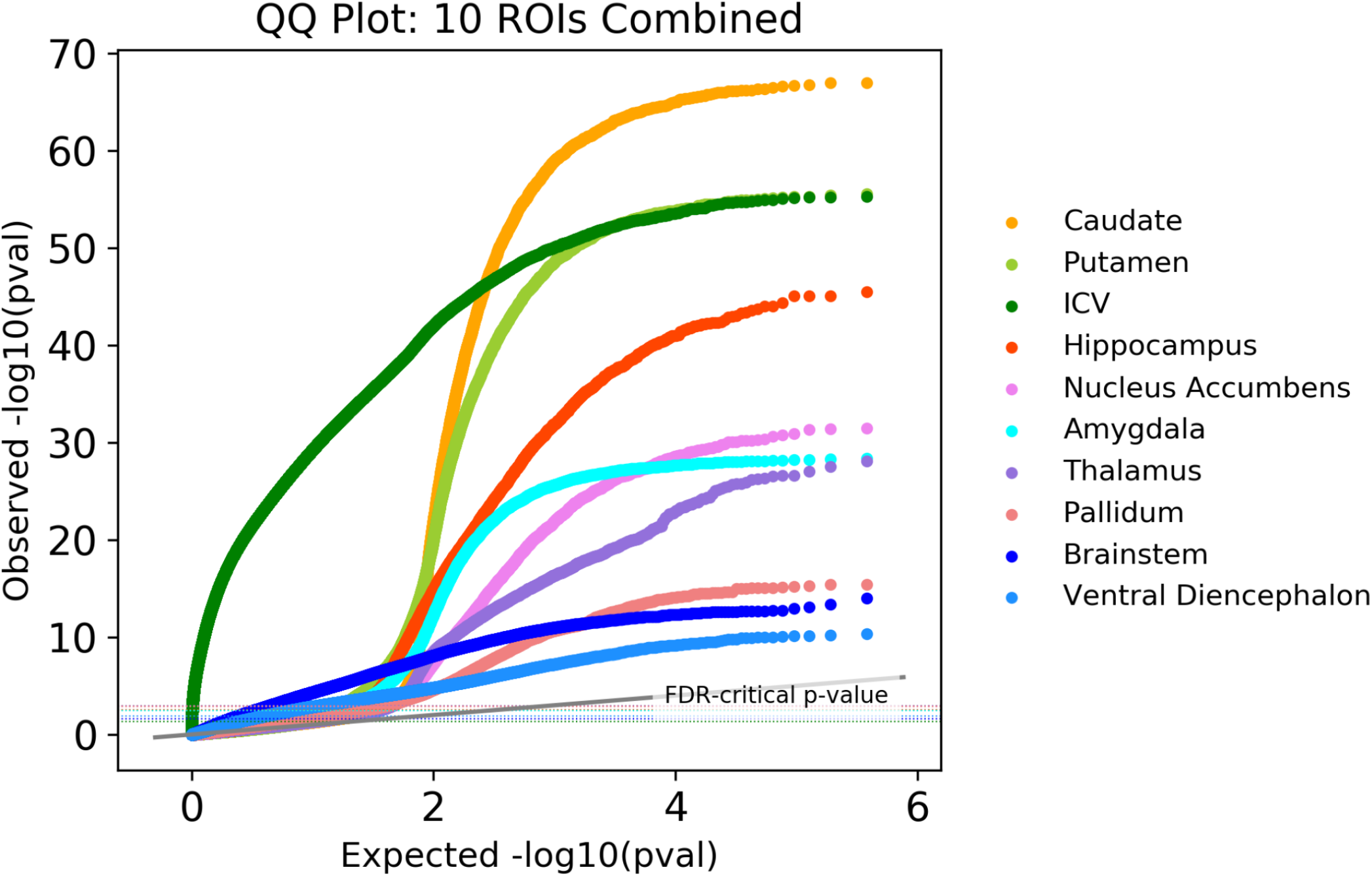
Quantile-Quantile (QQ) plots of raw *p*-values from all ten whole-brain voxel-wise linear mixed regression models for the association between polygenic score for brain ROI and GMV. FDR-correction (q=0.05) *p*-value thresholds are included as dashed lines for reference.

## III. Results and Discussion

Assessing the relationship between the polygenic score for each brain volume ROI and voxel-wise whole-brain volumes from this independent sample of 2,830 UKB participants, we observed that there was a statistically significant association with regional gray matter volume and each respective brain subvolume ROI PGS in each linear mixed regression model after correcting for multiple comparisons using the standard false discovery rate (FDR) at 5% (q=0.05). Voxel-wise whole-brain significance maps for the amygdala, hippocampus, putamen, thalamus, and the brainstem are shown overlaid on a brain map template (**Figure 2**: left → right, top → bottom). Their FDR-corrected significance values are reported in **Table 1** and summarized in **Figure 2** (caption).

Additionally, significant results were observed in the other ROIs: caudate nucleus (standard-FDR critical *p*-value=0.0012), nucleus accumbens (standard-FDR critical *p*-value=0.0014), globus pallidus (standard-FDR critical *p*-value=0.0013), ventral diencephalon (standard-FDR critical *p*-value=0.0128), as well as for ICV (run without TIV as a covariate to avoid overfitting; standard-FDR critical *p*-value=0.0498).

These results highlight the utility of VBM as a tool for mapping the effects of PGS across the whole brain, to determine whether these effects are general or regionally specific. By allowing for voxelwise statistical testing, VBM provides a spatially resolved framework to examine how aggregated genetic associations relate to gray matter volume beyond region-level summary measures. The PGS effect is often strongly, but not always, spatially localized to the anatomical boundaries of each subcortical ROI, as evidenced by those structures’ voxelwise significance in the association maps. Despite the polygenic nature of the traits under study, these effects are not diffusely distributed across the entire brain, but instead concentrate within the regions indexed by the corresponding PGS. For example, the statistical significance map for the linear mixed model analysis of the polygenic score for thalamus volume as the ROI and voxelwise whole-brain volumes shows the collective structure of the thalami clearly visible as significantly associated regions highlighted in our map after FDR-thresholding. The emergence of bilateral thalamic anatomy in these maps indicates that voxelwise associations form coherent, biologically meaningful spatial patterns rather than isolated effects. This shows that GWAS-derived PGS, while reflecting the cumulative influence of many genetic loci, give rise to anatomically interpretable and regionally specific patterns when mapped at the voxel level.

Further, our QQ plots of statistical significance values from each of the regression analyses in **Figure 3** show that larger ROIs typically yield similar or greater statistical significance than smaller ROIs; the curves show pronounced, region-specific departures from the null distribution (diagonal). Striatal regions, including the caudate nucleus and putamen, show the strongest and earliest deviations from the null, consistent with dense and spatially distributed genetic effects. Conversely, the hippocampus, nucleus accumbens, thalamus, and amygdala results display attenuated deviations from the null. Additionally, the larger subcortical ROIs, which included ventral diencephalon and brainstem, exhibit more shallow deviations.

Using QQ plots as an overall measure of the strength of association, these *p*-value distributional patterns are in line with the FDR-corrected results in **Table 1**, with ROIs showing smaller corrected *p*-values generally corresponding to curves that rise well above their ROI-specific significance thresholds. Overall, these findings address the central question of whether genetic influences identified through conventional GWAS act locally or more broadly across the brain. For subcortical brain volumes, we found that polygenic effects are primarily localized to their target structures– providing a spatially resolved link between GWAS discoveries and brain organization– and further demonstrating that voxelwise PGS mapping complements multivariate imaging genetics by revealing how distributed genetic architecture manifests in specific, anatomically interpretable neuroanatomical patterns.

## IV. Conclusions and Future Directions

In this independent sample of 2,830 participants from the UKB, we identified highly significant, localized effects for polygenic scores (PGS) associated with the amygdala, thalamus, and components of the basal ganglia. In contrast, PGS for brainstem volume showed widespread associations across the brain, rather than region-specific effects. These spatially distinct and overlapping brain-wide maps reveal both modular and distributed patterns of genetic influence on brain structure. Together, these findings provide a new perspective on the genomic architecture underlying individual variation in brain morphology.

The overall construction of this paper suggests a future direction where encoder-decoder transformers [37] are adapted to support the discovery of associations between arbitrary functions of the genome and arbitrary functions of brain images [18]. Our ability to identify connections between features in large-scale genomic and imaging data is likely to be enhanced by transformers whose encoder-decoder architecture uses cross-modal attention. In these approaches, optimal projections of the genome and the image are learned, to optimize mutual information between the projections, or predictive loss. Recent work on multimodal vision-language models (VLMs) suggests this connection could be achieved, once sufficient data are available, via CLIP-based projections or patch-based projections [38,39]. We are currently working on centralizing enough data to train these cross-modal models, and understanding how much data is sufficient to identify accurate and robust cross-modal associations [40].

## Acknowledgments

This research was supported by NIA grants R01AG058854 (ENIGMA World Aging Center), U01AG068057 (AI4AD), and an Alzheimer’s Association grant (to P.R.).

